# On the Evolution of Chromosomal Regions with High Gene Strand Bias in Bacteria

**DOI:** 10.1101/2024.02.28.582576

**Authors:** Jürgen Tomasch, Karel Kopejtka, Sahana Shivaramu, Izabela Mujakic, Michal Koblížek

## Abstract

On circular bacterial chromosomes, the majority of genes are coded on the leading strand. This gene strand bias (GSB) ranges from up to 85% in some Bacillota to little more than 50% in other phyla. The factors defining the extent of the GSB remain to be found. Here, we report that species in the phylum Gemmatimonadota share a unique chromosome architecture, distinct from neighboring phyla: In a conserved 600 kb region around the terminus of replication, almost all genes were located on the leading strands while on the remaining part of the chromosome the strand preference was more balanced. The high strand bias (HSB) region harbors the rRNA clusters, core, and highly expressed genes. Selective pressure for reduction of collisions with DNA replication to minimize detrimental mutations can explain the conservation of essential genes in this region. Repetitive and mobile elements are underrepresented, suggesting reduced recombination frequency by structural isolation from other parts of the chromosome. We propose that the HSB region forms a distinct chromosomal domain. Gemmatimonadota chromosomes evolved mainly by expansion through horizontal gene transfer and duplications outside of the ancient HSB region. In support of our hypothesis, we could further identify two Spiroplasma strains on a similar evolutionary path.

**Importance:** On bacterial chromosomes, a preferred location of genes on the leading strand has evolved to reduce conflicts between replication and transcription. Despite a vast body of research, the question why bacteria show large differences in their GSB is still not solved. The discovery of ‘hybrid’ chromosomes in different phyla, including Gemmatimonadota, in which a conserved high GSB is found exclusively in a region at *ter*, points towards a role of nucleoid structure, additional to replication, in the evolution of strand preferences. A fine-grained structural analysis of the ever-increasing number of available bacterial genomes could help to better understand the forces that shape the sequential and spatial organization of the cell’s information content.

## Introduction

Most bacterial chromosomes are circular with replication starting at one origin (*ori*) and progressing in both directions towards the terminus (*ter*). Since the earliest completely sequenced genomes, it became apparent that the need for an efficient integration of replication and transcription dictates the chromosome structure [1–3]. For example, highly expressed genes tend to be located closer to *ori*, taking advantage of remaining longer in a duplicated state while the DNA is copied. This is in particular the case for rRNA gene clusters that make up for 90% of bacterial RNA content [4]. Another constraint on gene arrangement is the possibility of clashes between the replication and transcription machineries as they move with high speed along the chromosome [5]. Both work with 5’-3’-directionality. The DNA Polymerase copies the leading and lagging strand in the direction of, and opposing the progressing replication fork, respectively. Frontal collisions between DNA and RNA Polymerase complexes slow down the transcription of lagging strand genes and can cause detrimental mutations [6,7]. Indeed, a preferential encoding of genes on the leading strand – co-directional to replication – seems to be the rule for bacterial chromosomes and it has been controversially discussed, how genes prevail on the lagging strand despite the accompanying negative effects [8–13].

There are large differences in the extent of the observed gene strand bias (GSB) between bacterial phyla [14–17]. In particular, on chromosomes of Bacillota (synonym Firmicutes) often more than 75% and up to 85% of all genes are encoded on the leading strand, while in most other phyla the distribution of genes between both strands is more balanced. To date, no satisfactory explanation has been found for these differences. In few bacterial phyla, including the Bacillota, the leading and lagging strand are replicated by utilization of two distinct Polymerase subunits, PolC and DnaE respectively, while in all others DnaE is responsible for replication of both strands [18]. It has been suggested that PolC activity might be responsible for maintenance of a strong GSB [19]. However, this hypothesis was not supported when a wider range of genomes from PolC-positive and -negative phyla were analyzed [16].

The phylum Gemmatimonadota comprises currently of only six cultured representatives. However, their ecological importance is underpinned by the discovery of hundreds of metagenome-assembled genomes (MAGs) from diverse environments [20–22]. Here, we report that the chromosome of our model strain *Gemmatimonas (Gem.) phototrophica* AP64 [23] contains a region near *ter* with an exceptional high GSB comparable to the Bacillota, while in the remaining part, genes showed a rather low preference for the leading strand. This high strand bias (HSB) region was also conserved in the other four Gemmatimonadota isolates with complete genomes. We further analyzed various PolC-positive and -negative bacterial phyla to assess the occurrence of similar chromosome architectures. We aimed to clarify how a clustered GSB can emerge and what could explain its evolutionary stability.

## Results

### Quantitative assessment of the gene strand bias

In order to identify regions with strong GSB, we chose an approach developed by de Carvalho and Ferreira [24]. The cumulative strand bias is calculated by moving along the chromosome of an organism and adding +1 for each gene on the plus and -1 for each gene on the minus-strand. As exemplified for *Bacillus subtilis* and *Escherichia coli* as representatives with a high and low strand-bias, respectively (Figure 1A), This approach results in curves with a steep and flat slope, respectively, positive for the right and negative for the left replichore (Figure 1B). Next, the cumulative GSB is correlated with the ascending positions of the respective genes for sliding windows along the chromosome. If all genes are positioned on the plus or minus strand, a correlation of 1 or -1 will be the result, respectively. For a random distribution, a value closer to 0 would be expected (Figure 1C). In the following analysis we use the squared correlation, correcting for the direction of the bias, referred to as the strand bias score (SBS).

**Figure 1:**
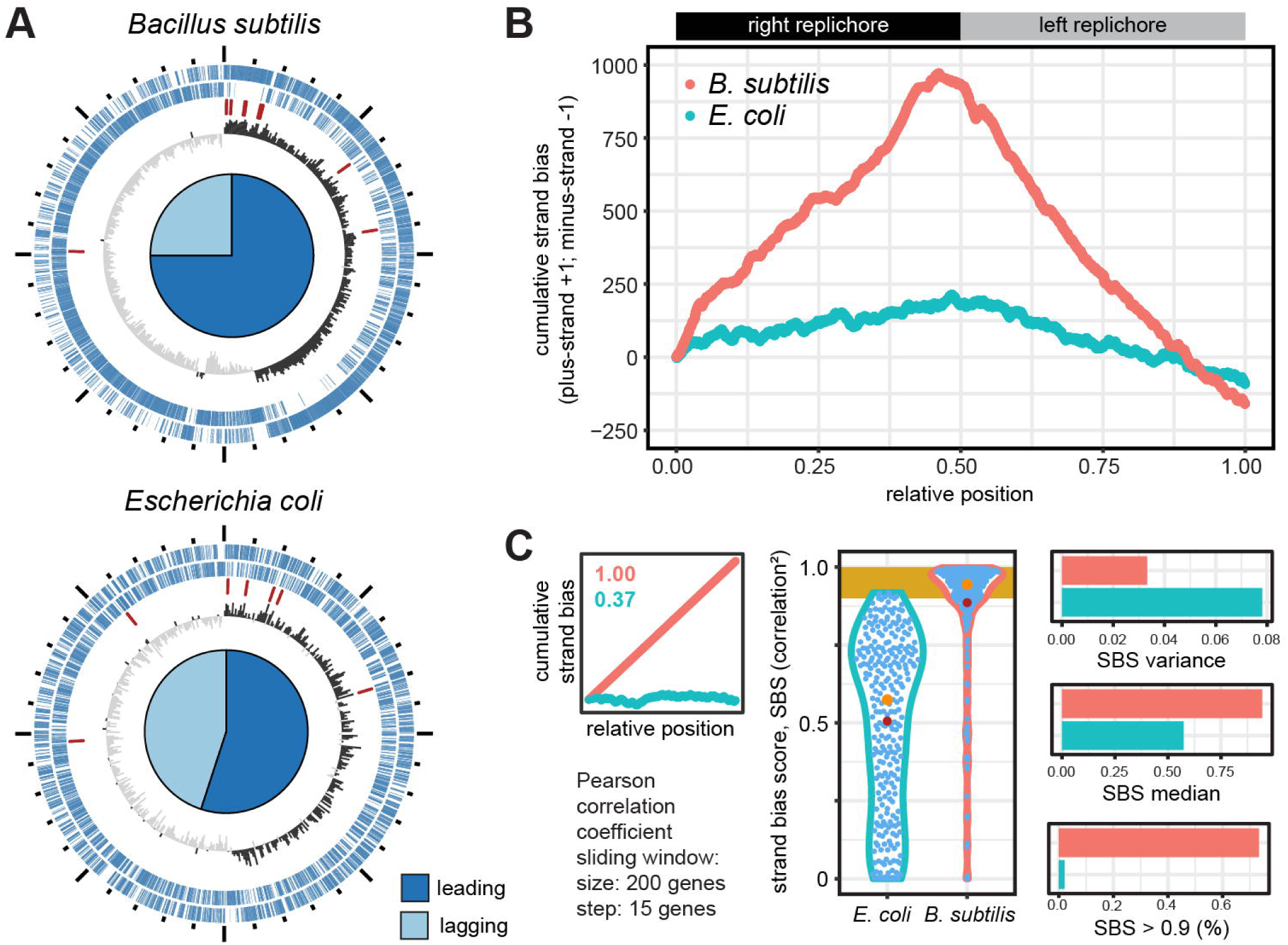
Analysis of the strand bias on circular bacterial chromosomes. **(A)** Chromosome structures of the model organisms *Bacillus subtilis* and *Escherichia coli*, with high and low strand bias, respectively. Depicted, from outer to inner ring, are protein-coding genes on the plus and the minus strand, rRNA genes and the GC skew. The plots are oriented with the origin of replication at the top. The proportion of genes on the leading and lagging strand is depicted as pie chart. **(B)** Cumulative strand bias for both organisms. For each gene on the plus and minus strand +1 and -1 are added, respectively. Counting starts on the right replichore. The position of the genes is normalized to chromosome size. **(C)** Strategy for identification of chromosomes with high and low strand bias. The strand bias score (SBS) calculated as squared correlation of the cumulative strand bias with gene position, for sliding windows of 200, moving by 15 genes (left panel). Characteristics of the distribution of the SBS (middle panel) that are extracted are the median, variance, the percentage of sliding windows with SBS > 0.9 (right panel).

As we were interested in identifying larger sections of the chromosome with a conserved strand bias, we chose a sliding window size of 200 moving by 15 genes for each step. The distribution of all calculated SBS values will provide information about the overall chromosome structure. The two model organisms differ in the median and variance of the distribution of SBS values. Furthermore, the proportion of HSB regions, with an SBS higher than 0.9, is 78% for *B. subtilis* and close to zero for *E. coli* (Figure 1C). Characterization of the SBS distribution for all analyzed genomes can be found in Supplementary Table S1. For a bacterium with clustered GSB, as the Gemmatimonadota, we would expect both, a high variance of the SBS and a proportion of HSB regions higher than zero.

### Gene strand bias in Gemmatimonadota and related phyla

The phylum Gemmatimonadota branches early within the so-called Fibrobacteres, Chlorobi, and Bacteroidetes (FCB) group [25]. The closest earlier branching neighbors of this group with cultivated species are the Verrucomicrobiota and Planctomycetota (Figure 2A). The closest phylum within the FCB group are the Fibrobacterota. The chromosome of our model organism *Gem. phototrophica* is characterized by a region with conserved gene order shifting from the plus to the minus strand in a region around *ter*, as derived from *ori* prediction and the GC skew (Figure 2B). This HSB region also harbored the two rRNA gene clusters. The genome structure published in 2014 was additionally confirmed by long read sequencing (Supplementary Figure S1). The chromosomes of all five Gemmatimonadota strains showed a similar conservation pattern of the GSB, with a high variance but also a high proportion of regions with an SBS > 0.9. The cumulative GSB plot was characterized by steep increase and decrease around the *ter* (Figure 2C). The reminder of the chromosome showed regions with a weaker and also totally missing GSB.

**Figure 2:**
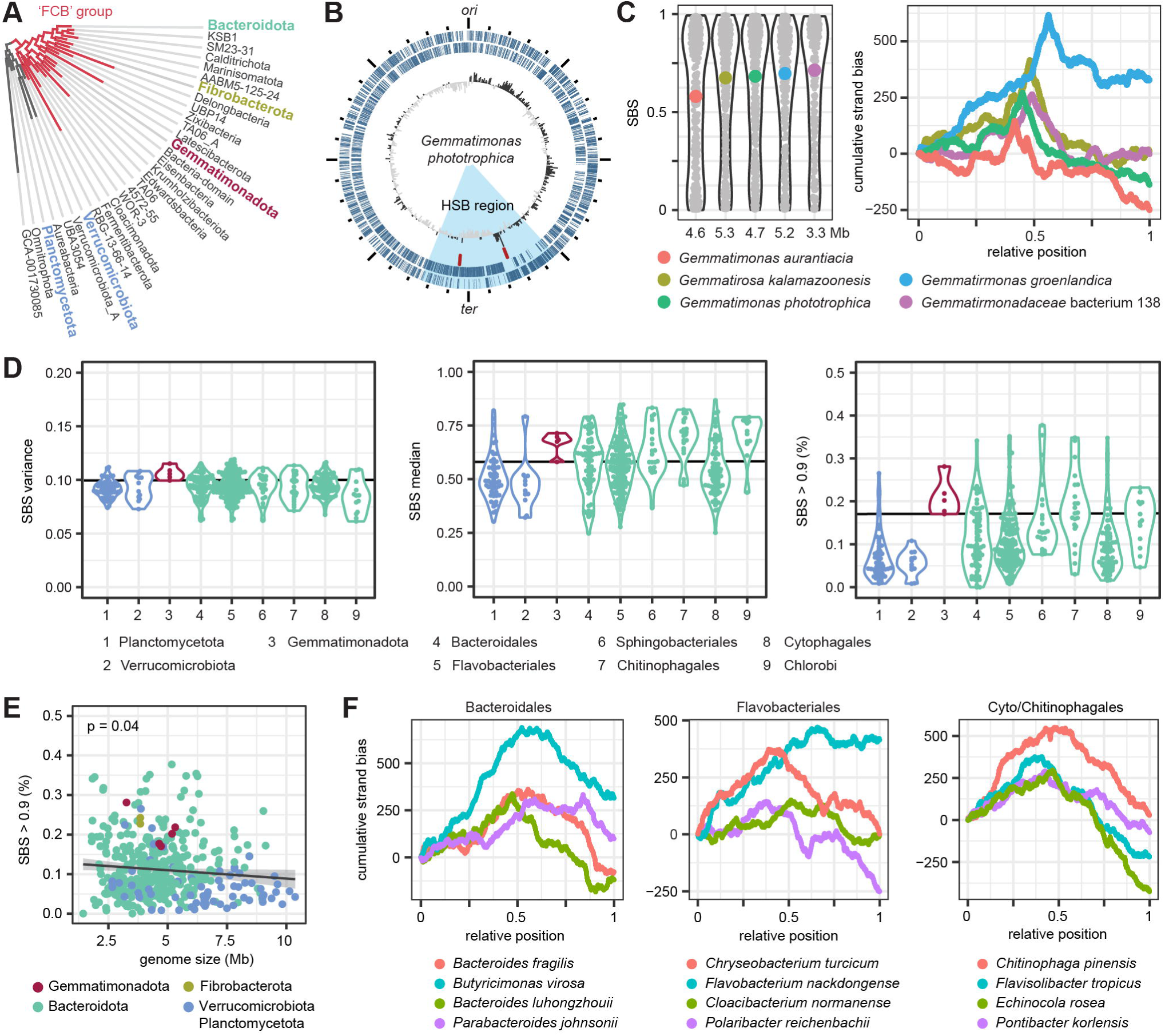
Strand bias in Gemmatimonadota and neighboring phyla. **(A)** Phylogenetic position of the Gemmatimonadota. Analyzed phyla are highlighted. The tree was obtained from GTDB using Annotree. **(B)** Chromosome of Gem. phototrophica AP64. The rings show genes on the plus and minus strand, the rRNA gene clusters and the GC skew. **(C)** Distribution of SBS and cumulative strand bias for the five Gemmatimonadota strains with closed genomes. **(D)** Variance and median of SBS in sliding windows, and the proportion of sliding windows with SBS > 0.9 in the phyla and orders colored as in Panel A. Data for the only two Fibrobacterota strains can be found in Supplementary Figure S2. **(E)** Relationship of correlation2 > 0.9 with genome size for all analyzed strains. The grey line indicates a fitted linear model. P-value of the slope is shown in the upper left corner. **(F)** Cumulative strand bias for chromosomes of selected Bacteroidota strains with high SBS variance and high number of segments with SBS > 0.9.

In an attempt to trace back the evolutionary origin of this chromosomal organization we searched for similar patterns in the neighboring phyla. The variance of the SBS was similar for all phyla but among the highest for the Gemmatimonadota who also showed a higher proportion of HSB regions relative to the others (Figure 2D). The chromosomes of the two earlier branching phyla, Verrucomicrobiota and Planctomycetota, were characterized by on average lower medians and, in particular, lower proportions of HSB regions than the FCB group strains. Within the Bacteroidota orders, the SBS medians and the number of HSB regions varied considerably. The latter ranged from almost none up to over 30% of the chromosome. Within the analyzed strains an increasing genome size was associated with a loss of conserved strand preference, in particular for the Verrucomicrobiota and Planctomycetota (Figure 2E).

For closer inspection, we selected Bacteroidota strains with an SDS variance (0.11) and proportion of HSB regions (0.17) at least as high as in the Gemmatimonadota (Figure 2F). A similar arrangement along the chromosome with the characteristic but less pronounced peak at *ter* was only found for *Bacteroides luhongzhouii* while the other strains showed a variety of patterns. For example, for *Butyricimonas virosa*, the typical V-shaped pattern of the cumulative strand bias indicated a moderate degeneration of gene orientation along the chromosome, but also showed two steep HSB stretches. In *Flavobacterium nackdongense* only the right replichore showed a strand bias. The different slopes for the left and right replichore in *Echinocola rosea* indicate a decay of the strand preference only on the latter.

One interesting case is *Fibrobacter succinogenes*, the closest relative to Gemmatimonadota among the analyzed strains. Here, a strong strand bias and stretches where it got lost are found in the *ori*- and *ter*-proximal half of the chromosome, respectively (Supplementary Figure S2).

In summary, the high variability of the strand bias between and within all analyzed phyla indicates a highly dynamic genome structure evolution. Strains with a conserved GSB along most of the chromosome were also found, although the degree of conservation was much lower than for previously reported Bacillota. The lack of clearly shared patterns makes it difficult to follow the evolutionary path of the Gemmatimonadota HSB region at this point.

### Features of strand biased regions compared to the rest of the Gemmatimonadota chromosomes

Next, we sought possible explanations for the emergence of the strand-biased region within the Gemmatimonadota. Therefore, we first analyzed the conservation of genes along the chromosome as a signature of genomic stability (Supplementary Table S2). As only five closed genomes from cultivated strains are currently available, we determined the pan-genome of the phylum by adding 61 previously curated, high quality MAGs [22]. Next, we analyzed the distribution of transposons (Supplementary Table S3) and repetitive DNA (Supplementary Table S4) on the chromosome. These factors potentially contribute to genomic dynamics and expansion [26]. The five genomes contained no complete and a maximum of two incomplete phages (Supplementary Table S5). These were not considered in the further analysis.

As exemplified for *Gem. phototrophica*, the HSB region differed in the studied characteristics from other parts of the chromosome (Figure 3A). In particular, the concentration of core genes and the absence of repeats became clearly apparent. For our model strain we also had transcriptome data available [27] and sought to identify differences in gene activity along the chromosome (Figure 3B). No replication-associated expression pattern was observed in accordance with the slow growth of the strain. Remarkably, the genes within the boundaries of the two rRNA operons, located inside the HSB region, showed a sharp increase in expression compared to the surrounding genes. While this region did not contain the most highly active genes, silenced and weakly expressed genes were completely absent. This points towards a physically separated cluster of high transcriptional activity at *ter*.

**Figure 3:**
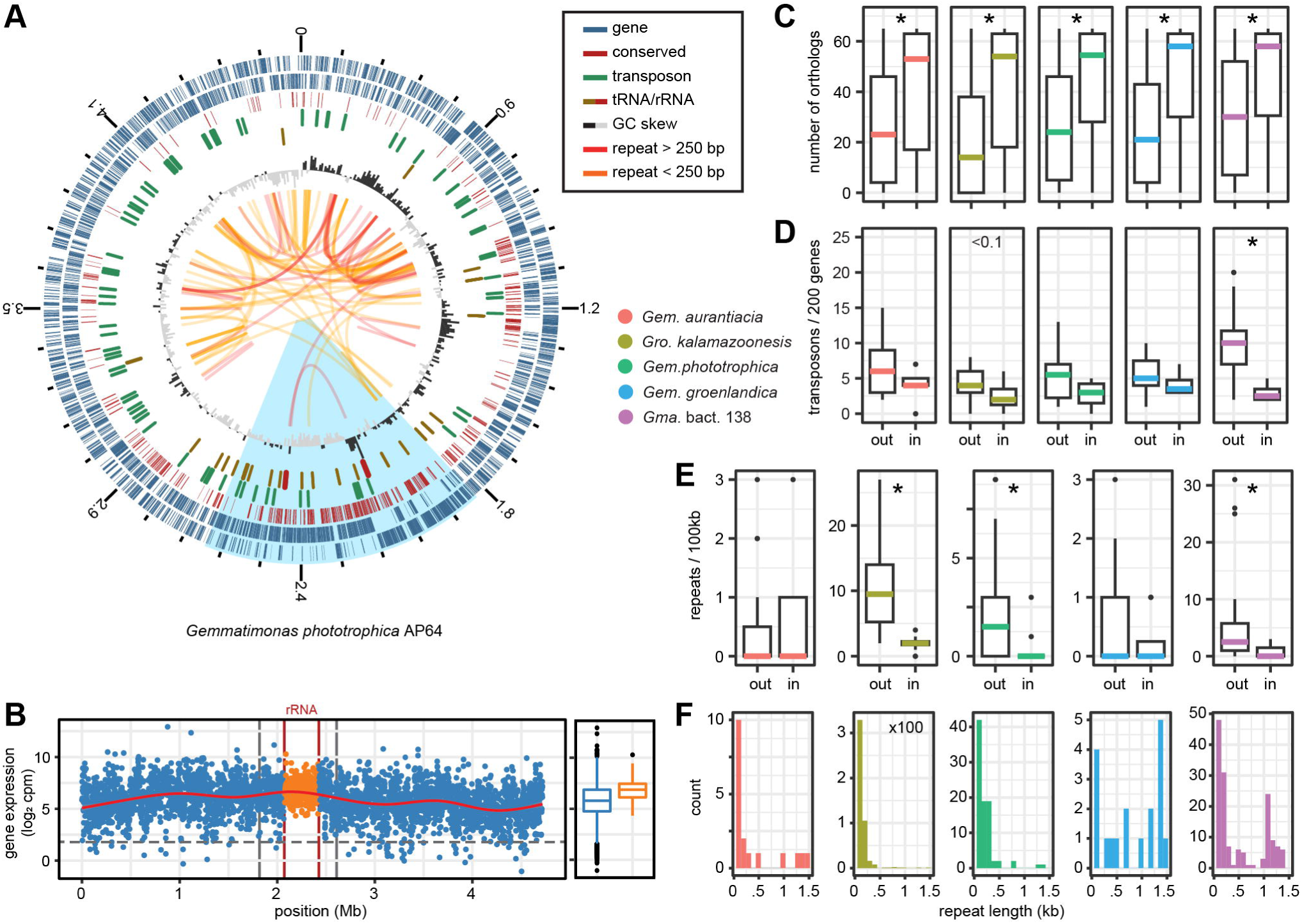
Chromosome structure of Gemmatimonadota. **(A)** Representative plot of Gem. phototrophica. The rings represent from outer to inner genes on the plus and minus strand, genes conserved in 61 out of 65 Gemmatimonadota genomes, tRNAs and rRNAs, transposons, and the GC skew. Small (80 to 250 bp) and large (>250 bp) repetitive elements are connected by yellow and red arcs, respectively. The plot is oriented with the origin of replication at the top. **(B)** Expression of genes along the chromosome in actively growing Gem. phototrophica as counts per million reads (cpm). The two rRNA operons and the region between them are indicated in red and yellow, respectively. Vertical gray lines mark the borders of the HSB region. **(C)** Comparison of the number of orthologues per gene, **(D)** transposons per 200 genes and **(E)** repetitive elements per 100 kb, outside and inside the strand biased region. **(F)** Distribution of repeat length identified in five Gemmatimonadota strains. Note the individual y-axis scale in (E) and (F) due to the large differences in numbers of repetitive elements between the strains.

In all five strains, conserved genes were significantly enriched in the HSB region (Wilcoxon test, p < 0.05). The median number of orthologues per gene ranged from 53 to 58 for inside and 14 to 30 for outside this region (Figure 3C). The position of the rRNA operons inside the HSB region was also conserved in the other strains (Supplementary Figure S3). Among others, part of the ribosomal and tRNA/rRNA-modifying genes, as well as the NADH-Dehydrogenase operon and two clusters of cell division genes were found within this region (Supplementary Table S2). In *Gem. aurantiaca*, the density of core genes was visibly lower on the left replichore and the GSB region was shifted to the right replichore. The opposite arrangement was found in *Gem. groenlandica*.

The median number of transposable elements was always higher outside the GSB region (4-10 compared to 2-4 elements/200 genes). Due to the high variance of transposon distribution along the chromosomes, this difference was significant only for *Gemmatimonadota (Gma.)* bact. 138 (Wilcoxon test, p < 0.05) which had the smallest genome and the highest density of the respective genes (Figure 3D). The five classes of transposons with the highest numbers of copies per genome were found in all five strains (Supplementary Table S3). In particular, the ISArsp14 element was present in 22 to 47 copies per strain. Three transposon classes with higher copy numbers (7 to 11) were found exclusively in *Gma.* bact. 138. All strains had a smaller number of single copy transposon classes present.

The strains differed considerably in the number of repetitive elements found on the chromosome (Figure 3E). In *Gem. aurantiaca* only 19 and in *Gem. groenlandica* only 20 repeats were found, with no preference to the in- or outside of the HSB region. In all other strains the repeat density was significantly higher in the less strand-biased segments of the chromosome. *Gem. phototrophica* had 86 repeats outside and only 4 repeats located inside the GSB region, two of the latter were the rRNA gene clusters. The *Gemmatirosa (Gro.). kalamazoonesis* chromosome was particularly densely packed with on average 11 copies/100 kb of two classes of short repeats, 90 and 150 bp in size and only few longer elements (471 in total). They were also found within the HSB region but with reduced density (Supplementary Figure S3). The chromosome of *Gma.* bact. 138 showed the greatest diversity of repeats. It shared the 150 bp short sequences with *Gro. kalamazoonesis*, but was also rich in 1050 bp long repeats found in clusters exclusively outside of the HSB region (157 in total).

In summary, the concentration of core genes and absence of repeats indicate that the Gemmatimonadota GSB region is ancient. We hypothesize that the parts of their chromosomes with a low GSB have evolved through genome expansion, either by horizontal gene transfer (HGT) or by duplication events.

### Clustered gene strand bias in bacteria with PolC DNA Polymerase subunit

To evaluate our hypothesis, we sought to identify such an expansion event in representatives of PolC-positive phyla, in which a high GSB is usually conserved along the chromosome. Besides Bacillota, forming the foundation for the proposed (and rejected) PolC-dependency of the GSB [19], homologs have been identified in Fusobacteriota, Mycoplasmatota (former Tenericutes) and Thermotogota [28]. In particular, the latter phylum has been previously found to lack a GSB [16]. The Bacillota showed overall the lowest variation and high median of the GSB (Figure 4A). For 116 out of 120 chromosomes the median SBS of the sliding windows was higher than 0.9. The PolC proteins of 12 strains had large alterations of the protein structure, for example losses of entire conserved domains (Supplementary Table S6). However, the strand bias was not reduced in any of these strains (Supplementary Figure S4). The other three phyla showed significantly different patterns (Tukey HSD, p-value < 0.05). The Fusobacteriota had retained overall a high strand bias although to a lesser extent and more variable than the Bacillota. The Thermatogota showed the highest variance and the lowest median and no more than 25% of the genes located in strand-biased regions. In these aspects their chromosomes resembled more that of PolC-negative *E. coli*. Genomes larger than 3 Mb, mostly present in Bacillota and Fusobacteriota, tended to have a higher gene strand bias (Figure 4B).

**Figure 4:**
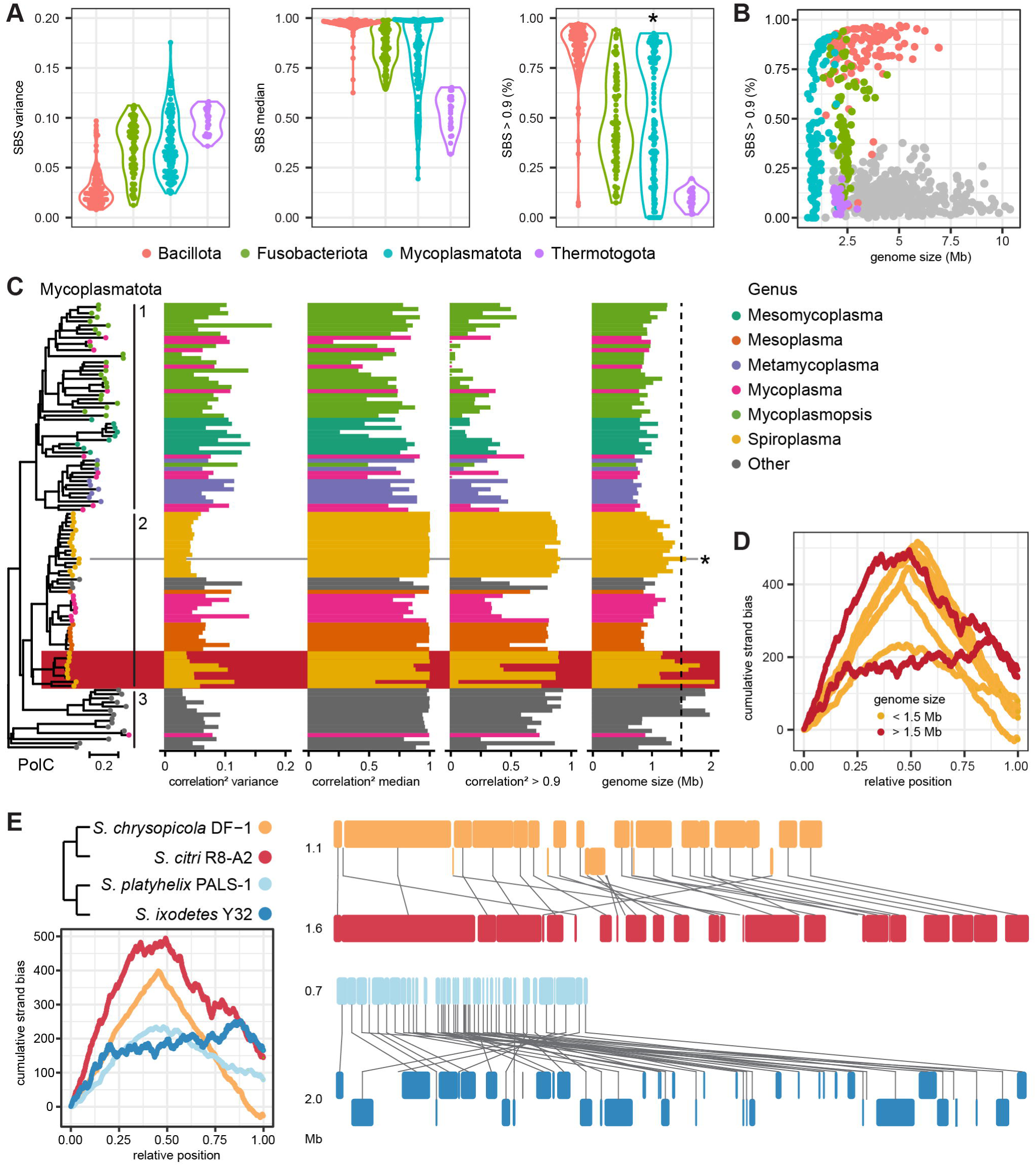
Strand bias in PolC-containing bacterial phyla. **(A)** Variance and median of SBS in sliding windows, and the proportion of sliding windows with SBS > 0.9 in the four phyla with *polC* encoded in the genome. The asterisk in the right panel indicates significant bimodality of the distribution. **(B)** Relationship of SBS > 0.9 proportion and genome size. Strains from Figure 2 are shown in grey for comparison. **(C)** Unrooted neighbor-joining tree of the Mycoplasmatota PolC protein compared to the three parameters of the chromosomal correlation^2^ distribution and genome size. Three distinct phylogenetic groups and an exceptional cluster of *Spiroplasma* strains are highlighted. A 1.5 Mb cut-off for a possible genome-expansion (dashed line) was set based on the previously published *S. clarkii* case (marked with an asterisk). **(D)** Cumulative strand bias for the chromosomes of the *Spiroplasma* strains highlighted in the previous panel. One potentially misassembled strain has been removed. **(E)** Cumulative strand bias and whole chromosome alignments of two pairs of closely related *Spiroplasma* strains.

Of all four phyla, the Mycoplasmatota showed the widest ranges of the analyzed parameters and a significant bimodality (excess mass test, p-value < 0.05) of the HSB proportion (Figure 4A). We investigated, therefore, if we could trace back the evolutionary loss of the GSB in specific genera within this phylum (Figure 4C). Genome reduction in the course of adaptation to an intracellular parasitic lifestyle is a key feature of Mycoplasmatota evolution [29].

Therefore, we defined a threshold of 1.5 Mb genome size for identification of potential expansion events. This value is slightly below the size of the *Spiroplasma (Spl.) clarkii* genome for which such an expansion through HGT has been documented, although without a change in the GSB [30]. Based on the PolC protein alignment, three different phylogenetic groups could be distinguished (Figure 4C). The first group, consisting mainly of *Meso*- and *Metamycoplasma* and *Mycoplasmopsis* had an overall low strand bias. Within the second group, a high strand bias was lost in *Mycoplasma* but conserved in most *Spiro*- and *Mesoplasma* genomes. The third group of genera with a low number of representative genomes showed both, low and high strand biases. In one *Spiroplasma* cluster, three exceptional large genomes showed a reduced proportion of segments with SBS > 0.9. In those, the strand bias was conserved near *ori*, but to a different extent lost towards *ter* (Figure 4D).

Comparing each to their closest relative, we observed two different evolutionary trends for the two pairs of strains (Figure 4E and Supplementary Figure S5). Large parts of the right replichore of the *Spl. citri* chromosome (1.6 Mb) showed a strong strand bias as present on the whole chromosome of its smaller relative *Spl. chrysopola* (1.1 Mb). However, in the *ter*- proximal region and parts of the left replichore, consisting of unique DNA, genes had no strand preference. *Spl. platyhelix* showed a deterioration of the strand bias along its small chromosome (0.7 Mb) as indicated by the reduced slope of the less pointy cumulative curve. Its relative *Spl. ixodetes* (2.0 Mb) maintained gene order only close to *ori*, while no strand preference was observed for the rest of the chromosome. The deviations between these two patterns could reflect independent gains, losses and inversions.

In summary, a high GSB was only conserved in the Bacillota but lost to different degrees in the other three PolC-positive phyla. Within the Mycoplasmatota, we could trace back the evolution of *Spl. citri* in which the ancient parts of the chromosome retained strand preference while newly acquired parts showed a less conserved GSB. This pattern is similar to our observations in Gemmatimonadota.

## Discussion

The dramatic increase in the number of sequenced bacteria in the last two decades has led to a broad understanding of the rules governing genome evolution. Examples are the surprisingly linear correlation between chromosomal GC-content and the C:N-ratio of the favorite carbon sources [31], or the preference for phage integration in proximity to *ter* [32]. A current study across 773 species, covering all major bacterial taxa, found conserved positions on the chromosome, in particular a bias towards *ori* and *ter*, for almost half of the identified gene families [33]. The *ori*- or *ter*-proximal position of regulatory genes can be strongly conserved across an order but can also show distinct evolutionary trajectories between phyla [34,35]. Exceptional genome architectures, defying the general evolutionary trend within a phylum, have also been found [30,36].

The questions of how the gene strand bias has emerged, is maintained and gets lost have been partially answered. Purifying selection can remove genes on the lagging strand if their expression interferes negatively with replication [8,12,37]. Contrastingly, the higher mutation rate might provide a fitness benefit for genes that need to be quickly adapted, like those coding for virulence or transcription factors [10,38]. Regardless of the detrimental effects, most bacteria thrive well with a rather large fraction up to almost half of their genes oriented head-on to replication [1,17,39]. Gene inversions, identified through a sign-change in GC-skew compared to the surrounding, seem to be common, although the frequency and directionality can vary between phyla [10,39]. The Bacillota remain so far the only exception with an almost universally conserved strong GSB, even along large chromosomes. The other PolC-positive phyla have diverged into clades with a different conservation of the GSB, like the Mycoplasmatota, or lost it completely like the Thermotogota. PolC might still be a necessary but is definitely not a sufficient prerequisite for a conservation of the strand bias along the full chromosome [16]. Thus, an explanation for the observed differences in the GSB is still lacking.

The discovery of bacteria with ‘hybrid’ chromosomes, having segments with both, high and low GSB, might help to understand the evolutionary development of strand preferences. The Gemmatimonadota chromosomes harbor a distinct 600 kb-region with a pronounced GSB switching from the plus to the minus strand. This region is roughly opposite of the *ori*, but the position of the sign switch does not always split the chromosome in equal halves. It has been shown before that the position of the *dif*-site, as a proxy for *ter*, relative to the *ori* can vary [40]. It might be possible that replication ends where genes switch their strand preference.

Consequently, the number of genes on the leading strand and, thereby, co-directionality of replication and transcription would be maximized. While opposing the general trend observed for other bacteria [4,41], the position of rRNA (and core) genes at the terminus of replication is characteristic for slow growing strains, in which gene dosage plays only a minor role [42]. Expression of these genes is presumably highest during cell division when new ribosomes and other important cell components have to be synthesized. The same holds true for the *ter*-proximal cell division genes that were found to be actively transcribed during replication in other bacteria [43,44]. Co-directional transcription close to *ter* would minimize collisions with the replication machinery. This chromosomal setup would ensure that highly expressed essential genes are shielded from accumulating mutations.

How can the evolution of the Gemmatimonadota chromosome structure be explained? We suggest the following scenario (Figure 5A): The chromosome of the last common ancestor (LCA) of the present strains was smaller than the 3.3 Mb of *Gma.* bacterium 138 and had already a *ter*-proximal ordered gene orientation. The clustering of core genes in the HSB region might be explained by both, newly acquired genes near *ori* and the loss of *ter*-proximal lagging-strand genes due to purifying selection [45]. An imbalance towards gene gain by HGT would increase the chromosome size [46]. Several transposon classes, present in all species, integrated into the LCA chromosome. From there, evolution of the strains took different paths. *Gma.* bact. 138 showed the least size expansion but integrated several unique transposons that have spread across the chromosome. *Gro. kalamazoonesis* on the other hand showed the highest number of repeats that might be partly responsible for the largest chromosome size of the analyzed strains. All three Gemmatimonas species had expanded genomes. Differences in the position of the HSB-region relative to *ori* between *Gem. aurantiaca* and *groenlandica* indicate different replichore preferences for integration of new DNA. Repetitive elements have spread in only three out of the five analyzed strains. They probably do not represent the primary cause of genome expansion, rather the consequence of individual events after the first acquisition of new genes (Figure 5B). They might still contribute to genome inversion and gene shuffling [26]. Of note, the repeat density within the HSB region was always lower than outside, indicating that this region is to some extent shielded from the invasive spread of repetitive elements that could partly explain the structural conservation of this region.

**Figure 5:**
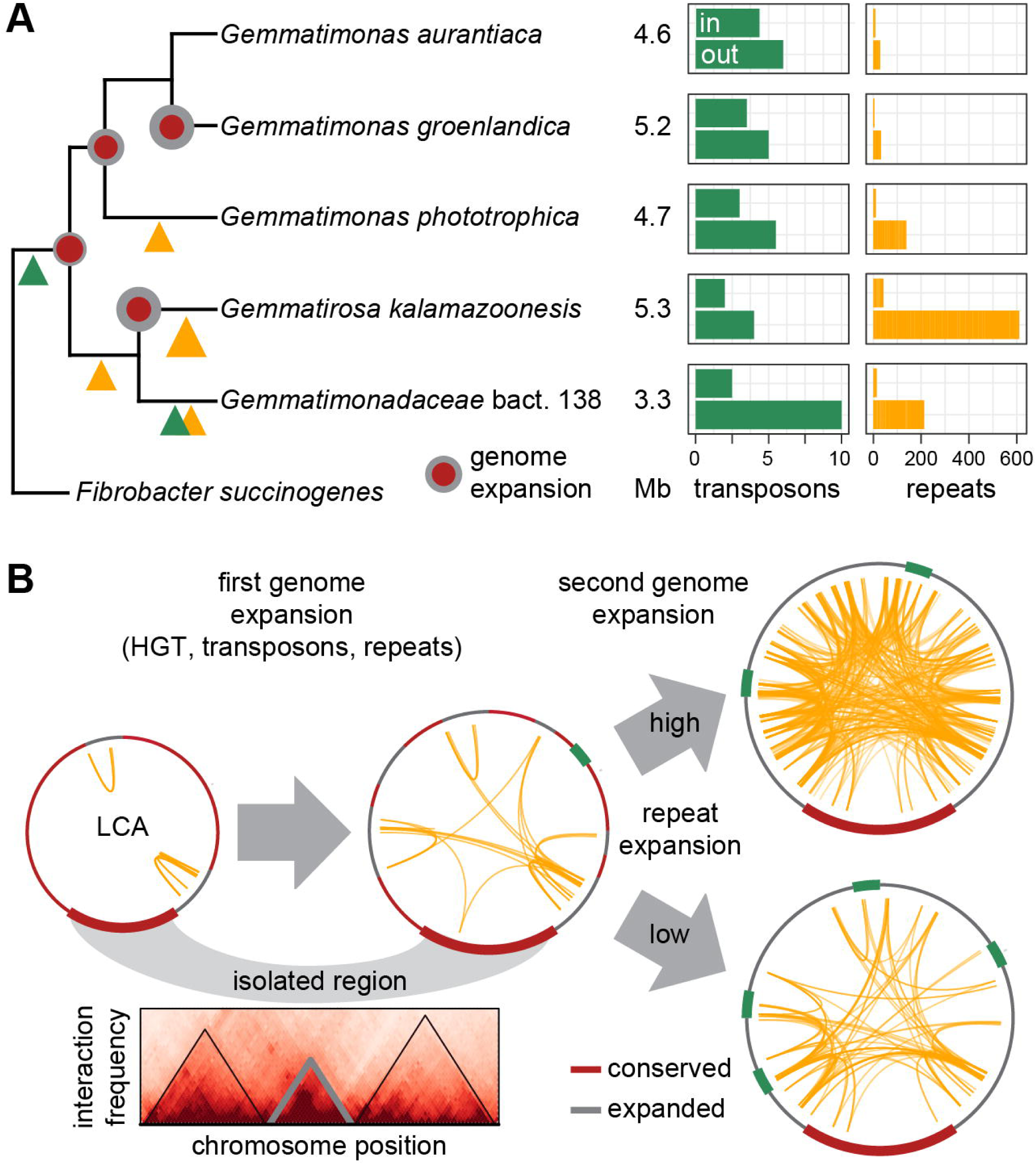
Evolution of the clustered gene strand bias (GSB) in Gemmatimonadota. **(A)** Cladogram of the analyzed strains based on 16S rRNA phylogeny. Major genome expansion events are indicated by circles at the nodes. Integration and spread of transposons and repetitive elements are indicated by triangles at the branches. The genome sizes in Mb are shown next to the strain names followed by the number of transposons per 200 genes and the total number of repeats inside and outside the GSB region. **(B)** Hypothetical scenario of Gemmatimonadota genome evolution with different timing and extent of chromosome restructuring events. Physical isolation could shield the GSB region on the folded chromosome from recombination and invasion of foreign elements. A theoretical chromosome interaction map illustrates this scenario. LCA, Last Common Ancestor.

To understand the evolutionary stability of the HSB region, it might be important to take the 3-dimensional chromosome structure into account (Figure 5B). Facilitated by several classes of structuring proteins, the chromosome folds into a highly condensed nucleoid in the cell. In this process subdomains are formed that can be isolated from each other [47–49].

Transcription further induces formation of boundaries between such domains [48,50]. The distinctively higher expression of genes between the two rRNA clusters could point to such a transcription-induced domain nested inside the Gemmatimonadota HSB region. A low interaction frequency with other DNA segments would reduce the probability of recombination and also the spread of mobile elements. We propose that the Gemmatimonadota HSB region forms an isolated chromosomal domain that allows coordination of transcriptional activity with replication, but also limits the contact to other parts of the chromosome. This hypothesis is in accordance with recent findings by Garmendina *et al.* who monitored recombination-coupled repair between two non-functional copies of a marker gene in Salmonella [51]. They showed that the probability for homologous recombination can vary greatly between individual chromosomal regions and is influenced by nucleoid-structuring proteins.

Our analysis was restricted to the Gemmatimonadota, their neighbors and PolC-positive phyla but nevertheless documents a highly dynamic evolution of the GSB and also revealed some unique gene distribution patterns. The ever-increasing number of available complete genome sequences will help to trace back the evolution of such remarkable chromosomal structures and help to understand the forces that shape the sequential and spatial organization of the cell’s information content.

## Material and Methods

### Dataset and tools

Genomes of the analyzed strains were obtained from the NCBI assembly database (accessed April 2023). Selection of phyla and families was guided by the GTDB database [52] and AnnoTree visualization [53]. We chose only type strains with a complete genome assembly. Only the chromosomes were considered and plasmids discarded.

Accession numbers can be found in Supplementary Table S1. For all Gemmatimonadota and for additional strains selected for visualization, chromosomes were centered around the *ori*, as determined by ori-finder website [54], using the reorientCircGenomes package in R [39]. Visualizations were realized using ggplot2 and ggbio [55]. A complete list of the programs used in the analysis can be found in Supplementary Table S7.

### Analysis of gene strand bias

The cumulative GSB was calculated as the sum of +1 for genes placed on the plus strand and -1 for gene on the minus strand. The squared correlation between the GSB and the chromosomal position, the SBS, was calculated for sliding windows of 200 kb moving in steps of 15 kb. The sliding window size was chosen to be one third of the Gemmatimonadota HSB region. Boundaries of the Gemmatimonadota HSB region were determined based on a step-wise increase in the SBS cut-off and manual curation. For each chromosome in the dataset, the SDS variance, median, and proportion of segments with SDS > 0.9 were calculated. In addition, the mean, kurtosis and skew of the distribution were calculated but not considered further in the analysis (Supplementary Table S1). Tukey’s HSD was used to compare the distributions of these parameters between strains from different phyla and families. The excess mass test was used to identify multimodal distributions [56]. Strains within the FCB group for which all three parameters were at least as high as the lowest value of the Gemmatimonadota were chosen for closer visual inspection.

### Analysis of genome conservation

For the Gemmatimonadota, the pan-genome was determined using proteinortho [57] using an e-value of 10^-15^ and 70% coverage as cut-offs. In addition to the five strains from ncbi we selected 64 MAGs from a previously analyzed dataset [22], with completeness > 90% and contamination < 5% as assessed by checkM [58]. For each gene of each strain we calculated the number of orthologs. Core genes were defined as being present in 90% of the genomes. Significant differences in the number of orthologs between chromosomal regions were identified with the Wilcoxon test. For the Mycoplasmatota, we selected the two strains with clustered strand bias and their closest relatives based on the PolC phylogeny and aligned each pair of chromosomes using mauve [59].

### Detection of mobile and repetitive elements

Transposable elements for each strain were retrieved by querying the ISfinder website with [60] with the protein coding genes and an e-value of 10^-5^ as cut-off. Only the best hit per gene was kept and the distribution of each class of insertion sequence was determined. Repeats were identified using repseek [61] which only accepts single entry fasta-files. A custom script was used to extract the largest sequence from the assembly genomic fasta file and pipe it directly into repseek with a minimal length of 32 bp for the initial seed. Repeats overlapping more than 80 bp (half the size of the smallest detected repeat) were counted as replicated entries and discarded. Based on visual inspection of repeat length distribution, two classes were assigned, shorter or longer than 250 bp. The Wilcoxon test was used to assess the significance of differences in transposon or read distribution between in- and outside of the GSB region. Phage DNA was identified using the Phaster website [62].

### Phylogenetic analysis

PolC amino acid sequences of Bacillota and Mycoplasmatota strains were retrieved from genomes downloaded from NCBI GenBank (Supplementary Table S1) Analyses were performed with MEGA 6.0 software [63]. Sequences were aligned using the ClustalW algorithm. Ambiguously aligned regions and gaps were manually excluded from further analysis. An unrooted phylogenetic tree was inferred by using the neighbor-joining algorithm with Jones-Taylor-Thornton model and 1,000 bootstrap replicates.

### Genome sequencing and analysis

Genomic DNA of *Gem. phototrophica* AP64 was extracted using the TIANamp Genomic DNA Kit (TIANGEN Biotech, Beijing, China). For obtaining high molecular weight genomic DNA, the CTAB method was used [64]. The complete genome was assembled by combining 150bp paired-end Illumina NovaSeq 6000 reads with Oxford Nanopore long-reads as described before [65].

### Data availability

Genomes of all analyzed strains are publically available at NCBI (https://www.ncbi.nlm.nih.gov/assembly). Accession numbers are provided in Supplementary Table S1. Scripts are available at github (https://github.com/Juergent79/gene_strand_bias).

## Acknowledgments

This work was supported by the Czech Science Foundation within the project PhotoGemm+ (GX19-28778X). The authors thank Alastair T. Gardiner for language correction.

## References

1. McLean MJ, Wolfe KH, Devine KM. Base Composition Skews, Replication Orientation, and Gene Orientation in 12 Prokaryote Genomes. J Mol Evol. 1998;47: 691–696. doi:10.1007/PL00006428

2. Rocha EPC. The replication-related organization of bacterial genomes. Microbiology. 2004;150: 1609–1627. doi:10.1099/mic.0.26974-0

3. Slager J, Veening J-W. Hard-Wired Control of Bacterial Processes by Chromosomal Gene Location. Trends in Microbiology. 2016;24: 788–800. doi:10.1016/j.tim.2016.06.003

4. Couturier E, Rocha EPC. Replication-associated gene dosage effects shape the genomes of fast-growing bacteria but only for transcription and translation genes. Molecular Microbiology. 2006;59: 1506–1518. doi:10.1111/j.1365-2958.2006.05046.x

5. Lang KS, Merrikh H. The Clash of Macromolecular Titans: Replication-Transcription Conflicts in Bacteria. Annual Review of Microbiology. 2018;72: 71–88. doi:10.1146/annurev-micro-090817-062514

6. Szczepanik D, Mackiewicz P, Kowalczuk M, Gierlik A, Nowicka A, Dudek MR, et al. Evolution Rates of Genes on Leading and Lagging DNA Strands. J Mol Evol. 2001;52: 426–433. doi:10.1007/s002390010172

7. Pomerantz RT, O’Donnell M. What happens when replication and transcription complexes collide? Cell Cycle. 2010;9: 2537–2543. doi:10.4161/cc.9.13.12122

8. Chen X, Zhang J. Why Are Genes Encoded on the Lagging Strand of the Bacterial Genome? Genome Biology and Evolution. 2013;5: 2436–2439. doi:10.1093/gbe/evt193

9. Gao N, Lu G, Lercher MJ, Chen W-H. Selection for energy efficiency drives strand-biased gene distribution in prokaryotes. Sci Rep. 2017;7: 10572. doi:10.1038/s41598-017-11159-3

10. Merrikh CN, Merrikh H. Gene inversion potentiates bacterial evolvability and virulence. Nat Commun. 2018;9: 4662. doi:10.1038/s41467-018-07110-3

11. Schroeder JW, Sankar TS, Wang JD, Simmons LA. The roles of replication-transcription conflict in mutagenesis and evolution of genome organization. PLOS Genetics. 2020;16: e1008987. doi:10.1371/journal.pgen.1008987

12. Liu H, Zhang J. Testing the adaptive hypothesis of lagging-strand encoding in bacterial genomes. Nat Commun. 2022;13: 2628. doi:10.1038/s41467-022-30000-8

13. Merrikh H, Merrikh C. Reply to: Testing the adaptive hypothesis of lagging-strand encoding in bacterial genomes. Nat Commun. 2022;13: 2627. doi:10.1038/s41467-022-30014-2

14. Rocha EPC, Danchin A. Ongoing Evolution of Strand Composition in Bacterial Genomes. Molecular Biology and Evolution. 2001;18: 1789–1799. doi:10.1093/oxfordjournals.molbev.a003966

15. Wu H, Qu H, Wan N, Zhang Z, Hu S, Yu J. Strand-biased Gene Distribution in Bacteria Is Related to both Horizontal Gene Transfer and Strand-biased Nucleotide Composition. Genomics, Proteomics & Bioinformatics. 2012;10: 186–196. doi:10.1016/j.gpb.2012.08.001

16. Saha SK, Goswami A, Dutta C. Association of purine asymmetry, strand-biased gene distribution and PolC within Firmicutes and beyond: a new appraisal. BMC Genomics. 2014;15: 430. doi:10.1186/1471-2164-15-430

17. Merrikh H. Spatial and Temporal Control of Evolution through Replication–Transcription Conflicts. Trends in Microbiology. 2017;25: 515–521. doi:10.1016/j.tim.2017.01.008

18. Dervyn E, Suski C, Daniel R, Bruand C, Chapuis J, Errington J, et al. Two Essential DNA Polymerases at the Bacterial Replication Fork. Science. 2001;294: 1716–1719. doi:10.1126/science.1066351

19. Rocha EPC. Is there a role for replication fork asymmetry in the distribution of genes in bacterial genomes? Trends in Microbiology. 2002;10: 393–395. doi:10.1016/S0966-842X(02)02420-4

20. Mujakić I, Piwosz K, Koblížek M. Phylum Gemmatimonadota and Its Role in the Environment. Microorganisms. 2022;10: 151. doi:10.3390/microorganisms10010151

21. Zheng X, Dai X, Zhu Y, Yang J, Jiang H, Dong H, et al. (Meta)Genomic Analysis Reveals Diverse Energy Conservation Strategies Employed by Globally Distributed Gemmatimonadota. mSystems. 2022;7: e00228–22. doi:10.1128/msystems.00228-22

22. Mujakić I, Cabello-Yeves PJ, Villena-Alemany C, Piwosz K, Rodriguez-Valera F, Picazo A, et al. Multi-environment ecogenomics analysis of the cosmopolitan phylum Gemmatimonadota. Microbiology Spectrum. 2023;0: e01112–23. doi:10.1128/spectrum.01112-23

23. Zeng Y, Feng F, Medová H, Dean J, Koblížek M. Functional type 2 photosynthetic reaction centers found in the rare bacterial phylum Gemmatimonadetes. PNAS. 2014;111: 7795–7800. doi:10.1073/pnas.1400295111

24. de Carvalho MO, Ferreira HB. Quantitative determination of gene strand bias in prokaryotic genomes. Genomics. 2007;90: 733–740. doi:10.1016/j.ygeno.2007.07.010

25. Gupta RS. The Phylogeny and Signature Sequences Characteristics of Fibrobacteres, Chlorobi, and Bacteroidetes. Critical Reviews in Microbiology. 2004;30: 123–143. doi:10.1080/10408410490435133

26. Rocha EPC. DNA repeats lead to the accelerated loss of gene order in bacteria. Trends in Genetics. 2003;19: 600–603. doi:10.1016/j.tig.2003.09.011

27. Shivaramu S, Tomasch J, Kopejtka K, Nupur, Saini MK, Bokhari SNH, et al. The Influence of Calcium on the Growth, Morphology and Gene Regulation in Gemmatimonas phototrophica. Microorganisms. 2023;11: 27. doi:10.3390/microorganisms11010027

28. Timinskas K, Balvočiūtė M, Timinskas A, Venclovas Č. Comprehensive analysis of DNA polymerase III α subunits and their homologs in bacterial genomes. Nucleic Acids Research. 2014;42: 1393–1413. doi:10.1093/nar/gkt900

29. Murray GGR, Charlesworth J, Miller EL, Casey MJ, Lloyd CT, Gottschalk M, et al. Genome Reduction Is Associated with Bacterial Pathogenicity across Different Scales of Temporal and Ecological Divergence. Molecular Biology and Evolution. 2021;38: 1570–1579. doi:10.1093/molbev/msaa323

30. Tsai Y-M, Chang A, Kuo C-H. Horizontal Gene Acquisitions Contributed to Genome Expansion in Insect-Symbiotic Spiroplasma clarkii. Genome Biology and Evolution. 2018;10: 1526–1532. doi:10.1093/gbe/evy113

31. Gralka M, Pollak S, Cordero OX. Genome content predicts the carbon catabolic preferences of heterotrophic bacteria. Nat Microbiol. 2023; 1–10. doi:10.1038/s41564-023-01458-z

32. Oliveira PH, Touchon M, Cury J, Rocha EPC. The chromosomal organization of horizontal gene transfer in bacteria. Nat Commun. 2017;8: 841. doi:10.1038/s41467-017-00808-w

33. Hu X-P, Lercher MJ. Nearly half of all bacterial gene families are biased toward specific chromosomal positions. bioRxiv; 2023. p. 2023.10.18.562889. doi:10.1101/2023.10.18.562889

34. Tomasch J, Koppenhöfer S, Lang AS. Connection Between Chromosomal Location and Function of CtrA Phosphorelay Genes in Alphaproteobacteria. Front Microbiol. 2021;12. doi:10.3389/fmicb.2021.662907

35. Koppenhöfer S, Lang AS. Patterns of abundance, chromosomal localization, and domain organization among c-di-GMP-metabolizing genes revealed by comparative genomics of five alphaproteobacterial orders. BMC Genomics. 2022;23: 834. doi:10.1186/s12864-022-09072-9

36. Kopejtka K, Lin Y, Jakubovičová M, Koblížek M, Tomasch J. Clustered Core- and Pan-Genome Content on Rhodobacteraceae Chromosomes. Genome Biology and Evolution. 2019;11: 2208– 2217. doi:10.1093/gbe/evz138

37. Zhang J, Yang J-R. Determinants of the rate of protein sequence evolution. Nat Rev Genet. 2015;16: 409–420. doi:10.1038/nrg3950

38. Mao X, Zhang H, Yin Y, Xu Y. The percentage of bacterial genes on leading versus lagging strands is influenced by multiple balancing forces. Nucleic Acids Research. 2012;40: 8210–8218. doi:10.1093/nar/gks605

39. Koppenhöfer S, Tomasch J, Lang AS. Shared properties of gene transfer agent and core genes revealed by comparative genomics of Alphaproteobacteria. Microbial Genomics. 2022;8: 000890. doi:10.1099/mgen.0.000890

40. Darling AE, Miklós I, Ragan MA. Dynamics of Genome Rearrangement in Bacterial Populations. PLOS Genetics. 2008;4: e1000128. doi:10.1371/journal.pgen.1000128

41. Rocha EPC, Danchin A. Gene essentiality determines chromosome organisation in bacteria. Nucleic Acids Research. 2003;31: 6570–6577. doi:10.1093/nar/gkg859

42. Vieira-Silva S, Rocha EPC. The Systemic Imprint of Growth and Its Uses in Ecological (Meta)Genomics. PLOS Genetics. 2010;6: e1000808. doi:10.1371/journal.pgen.1000808

43. Laub MT, McAdams HH, Feldblyum T, Fraser CM, Shapiro L. Global Analysis of the Genetic Network Controlling a Bacterial Cell Cycle. Science. 2000;290: 2144–2148. doi:10.1126/science.290.5499.2144

44. Alpers K, Vatareck E, Gröbe L, Müsken M, Scharfe M, Häussler S, et al. Transcriptome Dynamics of Pseudomonas aeruginosa during Transition from Overlapping To Non-Overlapping Cell Cycles. mSystems. 2023;8: e01130–22. doi:10.1128/msystems.01130-22

45. Fang G, Rocha EP, Danchin A. Persistence drives gene clustering in bacterial genomes. BMC Genomics. 2008;9: 4. doi:10.1186/1471-2164-9-4

46. Dagan T, Martin W. Ancestral genome sizes specify the minimum rate of lateral gene transfer during prokaryote evolution. Proceedings of the National Academy of Sciences. 2007;104: 870– 875. doi:10.1073/pnas.0606318104

47. Umbarger MA, Toro E, Wright MA, Porreca GJ, Baù D, Hong S-H, et al. The Three-Dimensional Architecture of a Bacterial Genome and Its Alteration by Genetic Perturbation. Molecular Cell. 2011;44: 252–264. doi:10.1016/j.molcel.2011.09.010

48. Cagliero C, Grand RS, Jones MB, Jin DJ, O’Sullivan JM. Genome conformation capture reveals that the Escherichia coli chromosome is organized by replication and transcription. Nucleic Acids Research. 2013;41: 6058–6071. doi:10.1093/nar/gkt325

49. Marbouty M, Le Gall A, Cattoni DI, Cournac A, Koh A, Fiche J-B, et al. Condensin- and Replication-Mediated Bacterial Chromosome Folding and Origin Condensation Revealed by Hi-C and Super-resolution Imaging. Molecular Cell. 2015;59: 588–602. doi:10.1016/j.molcel.2015.07.020

50. Bignaud A, Cockram C, Borde C, Groseille J, Allemand E, Thierry A, et al. Transcription-induced domains form the elementary constraining building blocks of bacterial chromosomes. Nat Struct Mol Biol. 2024; 1–9. doi:10.1038/s41594-023-01178-2

51. Garmendia E, Brandis G, Guy L, Cao S, Hughes D. Chromosomal Location Determines the Rate of Intrachromosomal Homologous Recombination in Salmonella. mBio. 2021;12: 10.1128/mbio.01151-21. doi:10.1128/mbio.01151-21

52. Parks DH, Chuvochina M, Rinke C, Mussig AJ, Chaumeil P-A, Hugenholtz P. GTDB: an ongoing census of bacterial and archaeal diversity through a phylogenetically consistent, rank normalized and complete genome-based taxonomy. Nucleic Acids Research. 2022;50: D785– D794. doi:10.1093/nar/gkab776

53. Mendler K, Chen H, Parks DH, Lobb B, Hug LA, Doxey AC. AnnoTree: visualization and exploration of a functionally annotated microbial tree of life. Nucleic Acids Research. 2019;47: 4442–4448. doi:10.1093/nar/gkz246

54. Dong M-J, Luo H, Gao F. Ori-Finder 2022: A Comprehensive Web Server for Prediction and Analysis of Bacterial Replication Origins. Genomics, Proteomics & Bioinformatics. 2022;20: 1207–1213. doi:10.1016/j.gpb.2022.10.002

55. Yin T, Cook D, Lawrence M. ggbio: an R package for extending the grammar of graphics for genomic data. Genome Biology. 2012;13: R77. doi:10.1186/gb-2012-13-8-r77

56. Ameijeiras-Alonso J, Crujeiras RM, Rodríguez-Casal A. Mode testing, critical bandwidth and excess mass. TEST. 2019;28: 900–919. doi:10.1007/s11749-018-0611-5

57. Lechner M, Findeiß S, Steiner L, Marz M, Stadler PF, Prohaska SJ. Proteinortho: Detection of (Co-)orthologs in large-scale analysis. BMC Bioinformatics. 2011;12: 124. doi:10.1186/1471-2105-12-124

58. Parks DH, Imelfort M, Skennerton CT, Hugenholtz P, Tyson GW. CheckM: assessing the quality of microbial genomes recovered from isolates, single cells, and metagenomes. Genome Res. 2015;25: 1043–1055. doi:10.1101/gr.186072.114

59. Darling AE, Mau B, Perna NT. progressiveMauve: Multiple Genome Alignment with Gene Gain, Loss and Rearrangement. PLOS ONE. 2010;5: e11147. doi:10.1371/journal.pone.0011147

60. Siguier P, Perochon J, Lestrade L, Mahillon J, Chandler M. ISfinder: the reference centre for bacterial insertion sequences. Nucleic Acids Research. 2006;34: D32–D36. doi:10.1093/nar/gkj014

61. Achaz G, Boyer F, Rocha EPC, Viari A, Coissac E. Repseek, a tool to retrieve approximate repeats from large DNA sequences. Bioinformatics. 2007;23: 119–121. doi:10.1093/bioinformatics/btl519

62. Arndt D, Grant JR, Marcu A, Sajed T, Pon A, Liang Y, et al. PHASTER: a better, faster version of the PHAST phage search tool. Nucleic Acids Research. 2016;44: W16–W21. doi:10.1093/nar/gkw387

63. Tamura K, Stecher G, Peterson D, Filipski A, Kumar S. MEGA6: Molecular Evolutionary Genetics Analysis Version 6.0. Mol Biol Evol. 2013;30: 2725–2729. doi:10.1093/molbev/mst197

64. Wilson K. Preparation of Genomic DNA from Bacteria. Current Protocols in Molecular Biology. 2001;56: 2.4.1–2.4.5. doi:10.1002/0471142727.mb0204s56

65. Neffe L, Abendroth L, Bautsch W, Häussler S, Tomasch J. High plasmidome diversity of extended-spectrum beta-lactam-resistant Escherichia coli isolates collected during one year in one community hospital. Genomics. 2022;114: 110368. doi:10.1016/j.ygeno.2022.110368

