## Supplementary Figures for "On the Evolution of Chromosomal Regions with High Gene Strand Bias in Bacteria"

This file contains: Supplementary Figures S1-S5

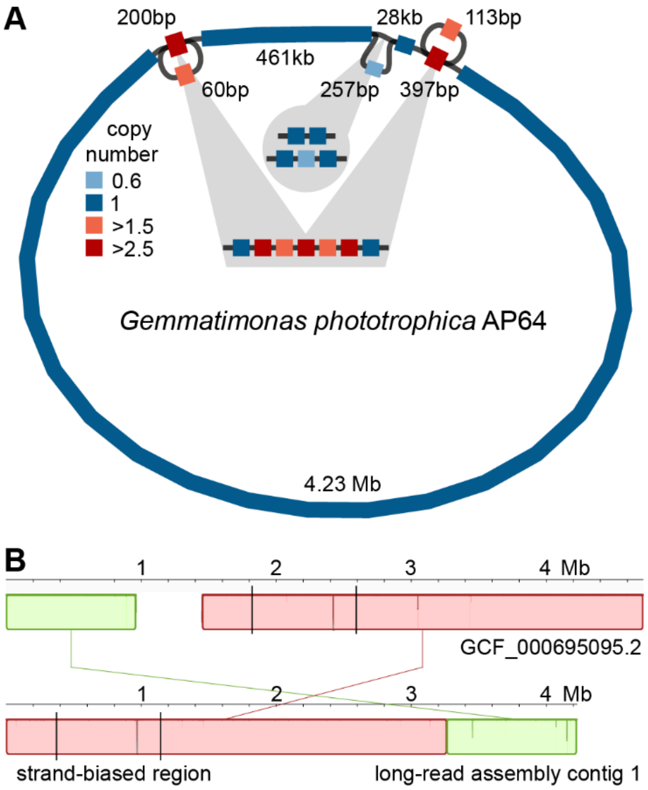


**Supplementary Figure S1 – Long-read resequencing of *Gemmatimonas phototrophica* AP64. (A)** Graph representation of the chromosome showing potential duplications and a loss of small DNA fragments. **(B)** Mauve whole genome alignment of the largest contig of the novel assembly to the reference strain. The strand-biased region is located approx. 190 kb from the contig end.

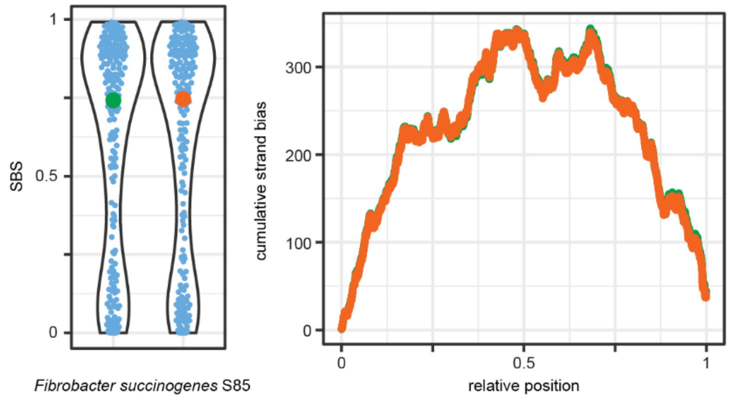


**Supplementary Figure S2 – Strand bias in two potentially clonal *Fibrobacter succinogenes* strains.** The distribution of correlation[2] and the cumulative strand bias along the chromosome is shown. In the latter small changes in the gene order become visible.

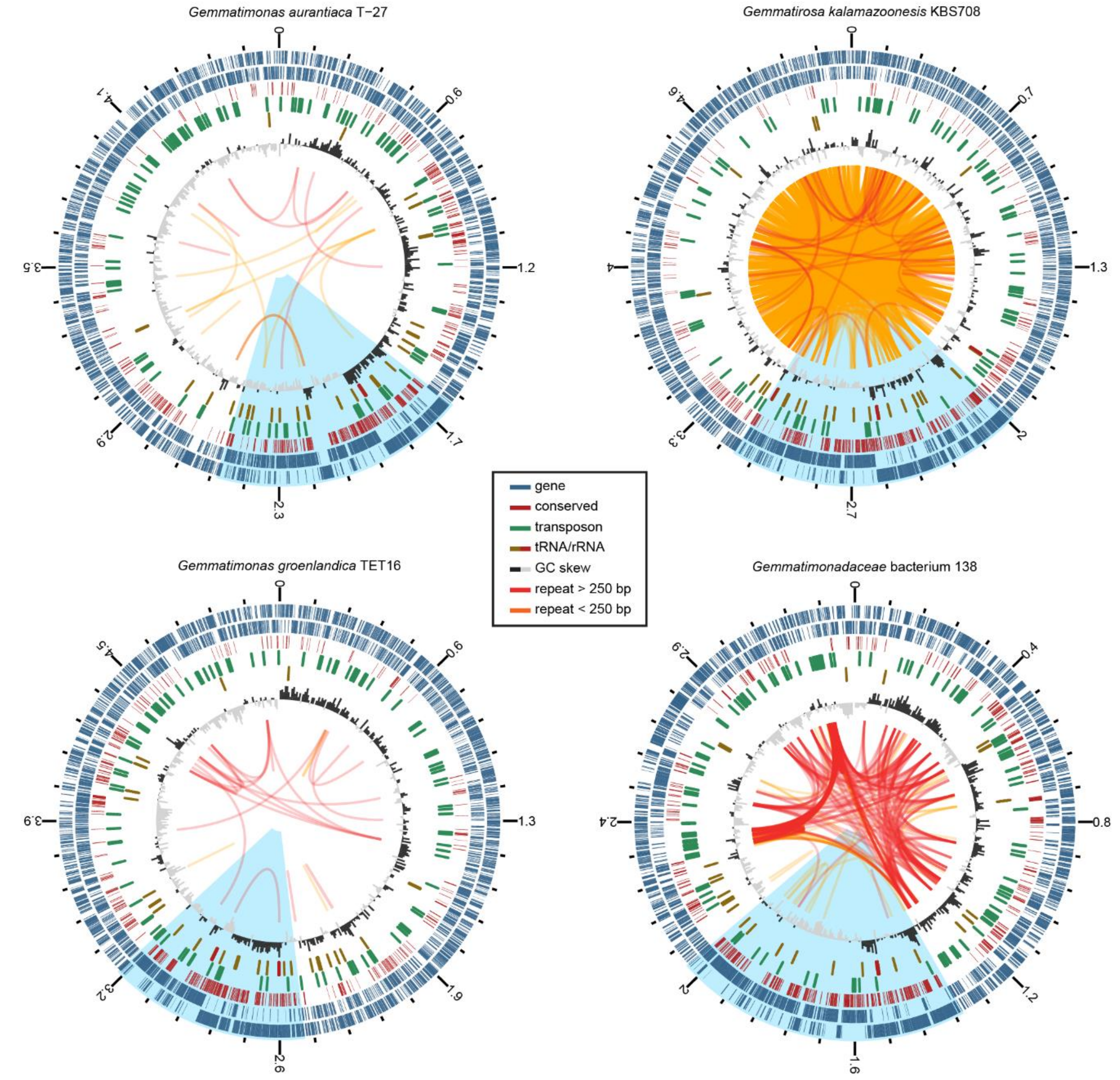


**Supplementary Figure S3 – Organization of Gemmatimonadota chromosomes.** The rings represent from outer to inner genes on the plus and minus strand, genes conserved in 61 out of 65 Gemmatimonadota genomes, tRNAs and rRNAs, transposons identified by IS finder (e-value < $10^{-15}$), and the GC skew. Small and large repetetive elements are connected by yellow and red arcs, respectively.

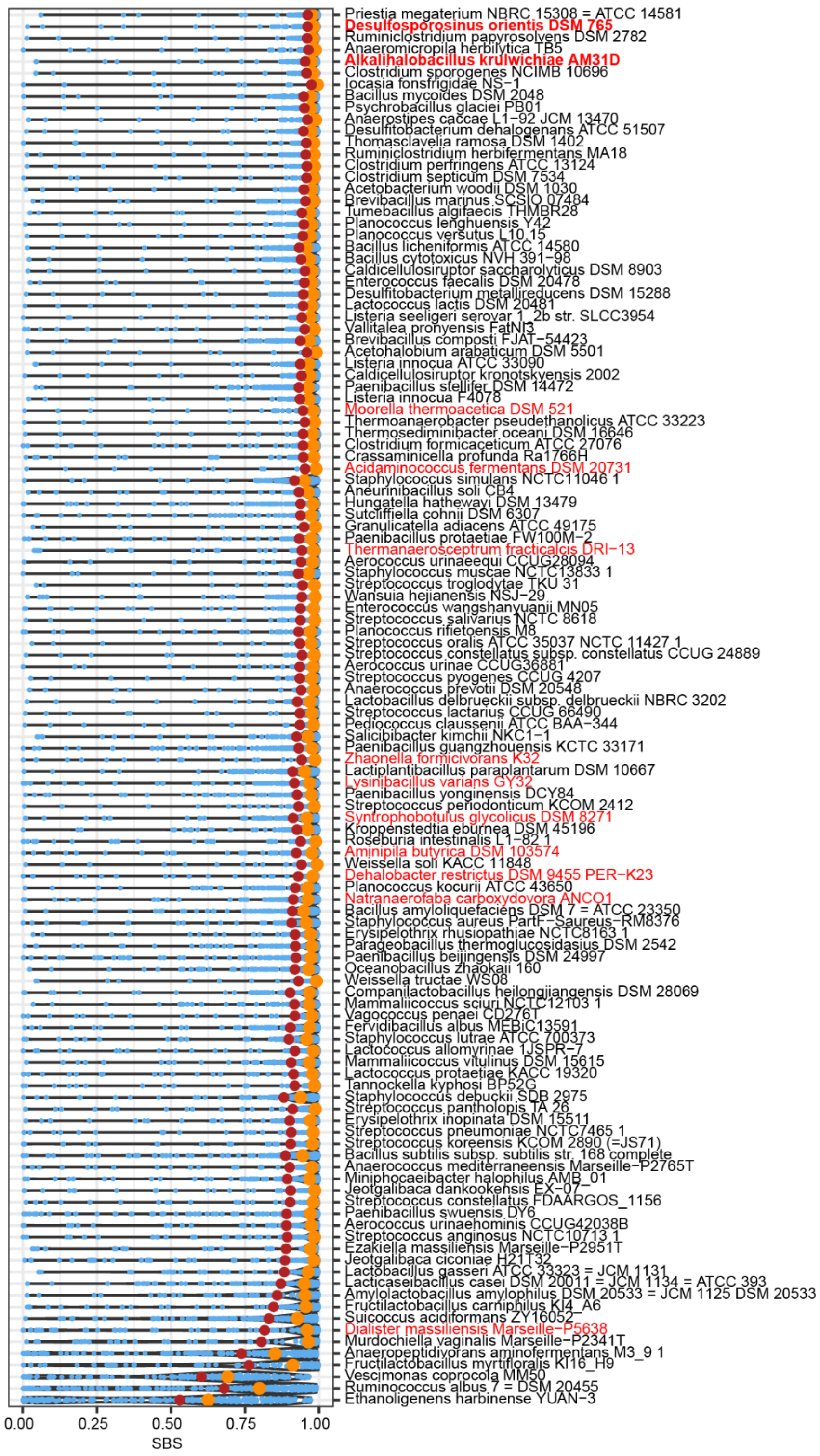


**Supplementary Figure S4 – Gene strand bias in Bacillota strains with alterations in PolC.** The highlighted strains showed large-scale insertion or deletions of the PolC sequence documented in Supplementary Table S6.

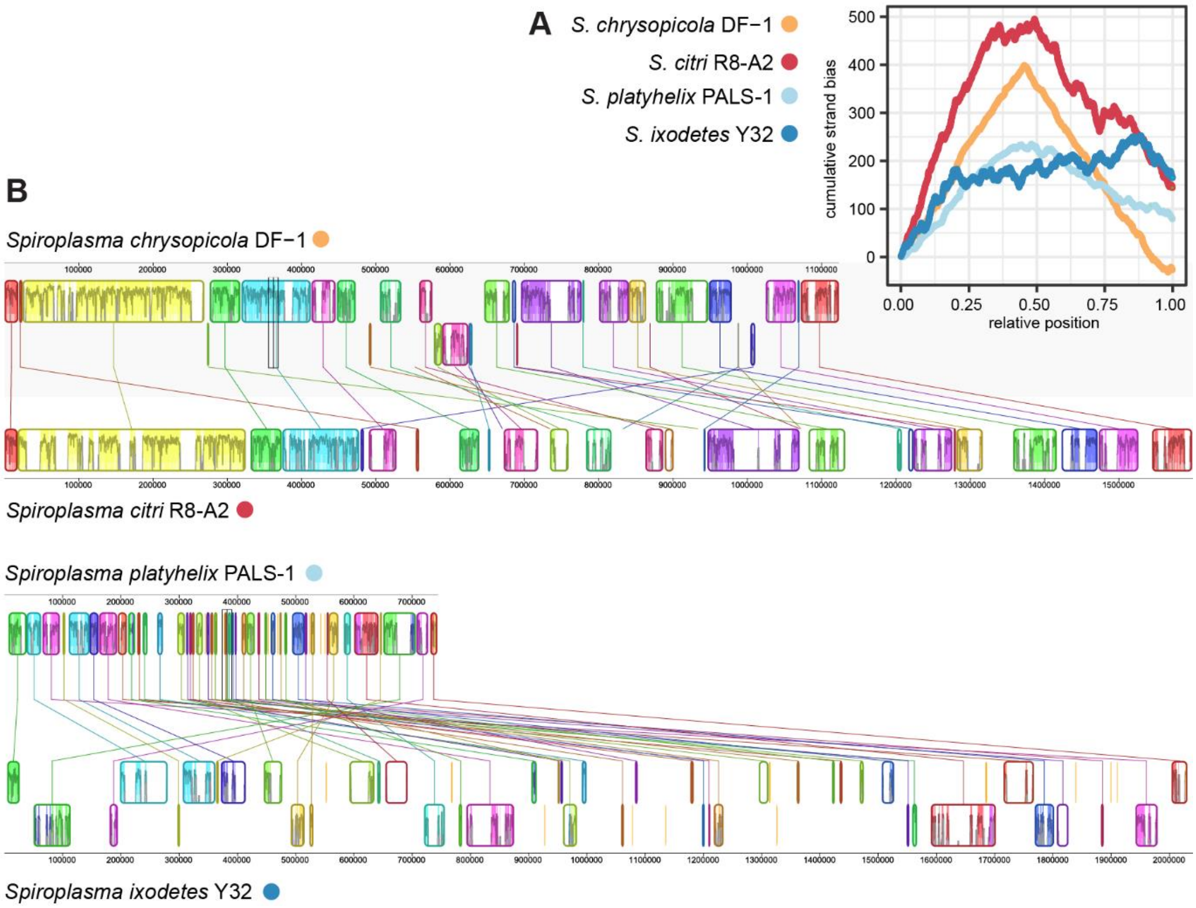


**Supplementary Figure S5 – Gene strand bias in *Spiroplasma* strains. (A)** Cumulative gene strand bias along the chromosomes of the two pairs of closely related Spiroplasma strains. **(B)** Mauve whole genome alignment of two pairs of closely related strains with different chromosome size and conservation of the gene strand bias.
